# A comparison between the neural correlates of laser and electric pain stimulation and their modulation by expectation

**DOI:** 10.1101/185645

**Authors:** EJ. Hird, D. Talmi, AKP. Jones, W. El-Deredy

## Abstract

**Background:** Pain is modulated by expectation. Event-related potential (ERP) studies of the influence of expectation on pain typically utilise laser heat stimulation to provide a controllable nociceptive-specific stimulus. Short painful electric stimulation has a number of practical advantages, but is less nociceptive-specific. We compared the modulation of electric versus laser-evoked pain by expectation, and their corresponding pain-evoked and anticipatory ERPs.

**New Method:** We developed understanding of recognised methods of laser and electric stimulation. We tested whether pain perception and neural activity induced by electric stimulation was modulated by expectation, whether this expectation elicited anticipatory neural correlates, and how these measures compared to those associated with laser stimulation. We elicited cue-evoked expectations of high and low pain and compared subjective ratings and corresponding ERPs in response to the delivery of laser and electric stimulation in a within-participant design.

**Results:** Despite sensory and affective differences between laser and electric pain, intensity ratings and pain-evoked potentials were modulated equivalently by expectation, though ERPs only correlated with pain ratings in the laser pain condition. Anticipatory correlates significantly differentiated pain intensity expectation to laser but not electric pain.

**Comparison with Existing Method:** Previous studies have consistently shown that laser-evoked potentials are modulated by expectation. We extend this by showing electric pain-evoked potentials are equally modulated by expectation, within the same participants. We also show a difference between the pain types in anticipation.

**Conclusions:** Though laser-evoked potentials express a stronger relationship with pain perception, both laser and electric stimulation may be used to study the modulation of pain-evoked potentials by expectation. Anticipatory-evoked potentials are elicited by both pain types, but they may reflect different processes and did not correlate with pain perception.

## 1. Introduction

Our experience of pain is profoundly influenced by what we expect to feel. Pain expectations can be experimentally manipulated through administration of a sham analgesic using a number experimental placebo procedures (Morton, Brown, Watson, El-Deredy, & Jones, 2010b; Wager, 2004; Watson et al., 2009a) or by eliciting cue-evoked expectations and testing the resultant pain report (Atlas, Bolger, Lindquist, & Wager, 2010; Brown, Seymour, Boyle, El-Deredy, & Jones, 2008a). The modulation of pain by expectation has received a significant amount of attention, which is reflected in recent meta-analyses and reviews (Amanzio, Benedetti, Porro, Palermo, & Cauda, 2013; Atlas & Wager, 2012; Brown et al., 2011; Finniss, Kaptchuk, Miller, & Benedetti, 2010; Jones & Brown, 2017; Petersen et al., 2014; Price, Finniss, & Benedetti, 2008). Expectations also change pain-related neural activity (Wager et al., 2004). However, the modulation of nociceptive responses to different modality stimuli by expectation has not been quantified.

ERPs^1^ are a useful tool to study nociceptive neural processing as they offer high temporal resolution and nociceptive specificity, allowing characterisation of instantaneous responses to pain. ERP studies have focused on the modulation of LEPs^1^ by expectation (Colloca et al., 2008; Lorenz et al., 2005; Lyby, Aslaksen, & Flaten, 2011; Martini, Lee, Valentini, & Iannetti, 2015; Morton, Brown, Watson, El-Deredy, & Jones, 2010b; Morton, Watson, El-Deredy, & Jones, 2009; Wager, Matre, & Casey, 2006; Watson et al., 2009b; Watson, El-Deredy, Vogt, & Jones, 2007). LEPs result from laser heat stimulation of nociceptive Aδ and c fibres (Iannetti et al., 2004; Iannetti, Zambreanu, & Tracey, 2006) and therefore have the advantage of being nociceptive-specific. LEPs are a well-validated method for assessing pain perception and its neural basis (Garcia-larrea, Frot, Valeriani, Bernard, & Lyon, 2003; Mobascher et al., 2009; Treede, Lorenz, & Baumgärtner, 2003).

An alternative method of pain induction is transcutaneous electrical stimulation, which activates myelinated Aβ somatosensory fibres as well as Aδ nociceptive fibres and elicits an EEP^1^. Across studies, it has been shown that both EEPs and LEPs express the P2 component which is closely linked to activity in the operculum, SII and the cingulate cortex (Bentley, Derbyshire, Youell, & Jones, 2003; Christmann, Koeppe, Braus, Ruf, & Flor, 2007; Garcia-larrea et al., 2003). Some studies have shown modulation of electrically induced pain by expectation, chiefly behavioural studies (Colloca, Sigaudo, & Benedetti, 2008; Luana Colloca & Benedetti, 2006, 2009b; Colloca, Petrovic, Wager, et al., 2010; De Pascalis, Chiaradia, & Carotenuto, 2002; Yeung, Colagiuri, Lovibond, et al., 2014), and fewer fMRI^1^ studies, showing modulation within the rACC^1^, insula and thalamus (Wager, 2004b), and ERP studies (Rütgen, Seidel, Riečanský, & Lamm, 2015). Laser and electric stimulation activate similar areas of the brain, sharing activation across key structures of the lateral and medial pain system. Expecting both types of stimulation modulates activity in areas such as the cingulate, insula, dorsolateral prefrontal cortices and thalamus (Amanzio et al., 2013; Bentley et al., 2003; Christmann et al., 2007; Wager, 2004b). As similar neural areas are activated by the two pain types, and as one would expect expectation to influence pain perception independent of the modality of pain stimulus, it is possible that expectation could modulate the ERP and intensity ratings of the two types of stimulation equivalently. We compare the effect of cue-evoked expectation on the P2 component of EEPs and LEPs.

Awareness of imminent pain elicits an anticipatory slow-wave EEG correlate termed the SPN^1^ (Böcker, Baas, Kenemans, & Verbaten, 2001). A marker for anticipation is a useful tool for quantifying the processes underlying expectation. The SPN has been characterised in response to laser pain as a negative potential peaking at central electrodes and has been localised to the cingulate and anterior insula, which is implicated in affective processing (Brown, Seymour, El-Deredy, & Jones, 2008a; Caria, Sitaram, Veit, Begliomini, & Birbaumer, 2010). The SPN in response to electric pain has been observed in posterior areas of the cortex (Berns et al., 2006; Hoflle, Pomper, Hauck, Engel, & Senkowski, 2013; Lin, Hsieh, Yeh, Lee, & Niddam, 2013), and centroparietal electrodes (Seidel et al., 2015); however, other studies have failed to show the SPN in electric compared to laser pain (Babiloni et al., 2003, 2007). Accordingly, the SPN to electric pain is yet to be reliably quantified.

The paucity of research into the modulation of electric pain by expectation may be because electric stimulation activates Aβ somatosensory fibres alongside Aδ nociceptive fibres which could add sensory noise to the signal (Perchet et al., 2012), activating a larger number of thalamo-cortical units than laser stimulation, and resulting in higher-amplitudes and EEPs compared to LEPs (Garcia-larrea et al., 2003; Gingold, Greenspan, & Apkarian, 1991; Treede, Kenshalo, Gracely, & Jones, 1999). This has likely led to the concern that the representation of somatosensory processes by the resultant EEP could interfere with measurement of expectation-induced pain modulation, as activity related to innocuous somatosensory activity could increase the noise of the EEP. Yet electrical stimulation has obvious advantages over laser stimulation, so it is important to understand whether we can study pain expectations with this technique. Laser stimulation can result in heating of the skin leading to sensitisation, which limits the number of trials in a study. Skin heating by the laser is also associated with the risk of skin lesions and therefore has additional ethical implications. The setup is expensive, may not be portable depending on the type of laser, requires operators to undertake substantial training, and requires the wearing of safety goggles which can be uncomfortable and distracting for participants. Transcutaneous electrical stimulation is arguably a more practical method of pain assessment. It does not heat the skin so there is no limit to the number of trials. There are fewer ethical implications in using electric stimulation to elicit pain, as the activation is transient and there is no risk of skin damage. Electric stimulators are cheaper, portable and available commercially, and require no specific training to use so can be used more widely and potentially in conjunction with physiological phenotyping in clinical trials to identify placebo responders. If EEPs are modulated by expectation equivalently to LEPs, this would allow generalisation across the literature, and future studies could use electric stimulation. We investigated ERP and the anticipatory SPN, and intensity ratings of laser and electric pain within the same participants, and predicted that they would be equally modulated by cue-evoked expectation.

## 2. Methods

### 2.1. Design

The study was a 2 (stimulator: laser/electric) × 2 (pain expectation: high/low) × 3 (pain intensity: high/low/medium) within-subjects design.

### 2.2. Participants

Twenty participants aged 18-35 (9 females, mean age 23 years) were recruited via newspaper and university advertisements and received £30 compensation for participation in the study. Participants had normal or corrected-to-normal vision. They had no history of neurological or psychiatric conditions, did not take medications which could affect their neurotransmitter levels, or take analgesics, and did not have a history of chronic pain. Ethical approval was granted by the University of Manchester, where the study took place.

### 2.3. Apparatus

Visual stimuli were presented on a desktop computer screen 1 metre from the participant. The laser stimuli were generated by a class 4 thulium laser (IPG Photonics Corporation, US/ TLR-30-2050, wavelength 2050nm/ 30 watt). The laser stimuli were of 150ms duration and a beam diameter of 5 millimetres. Energy delivered at 50% intensity was 2.25 joules. See table 1 for a summary of the energy densities of the laser stimulator. The electric stimuli were delivered by two silver/silver chloride cup electrodes attached to a TENS machine (maximum voltage 477 Volts; maximum current 81 milliamps) (Medical Physics, Salford Royal NHS Foundation Trust) and were of 230 to 300 microsecond duration. Maximum voltage was 477 volts; maximum current was 81 milliamps, and maximum output power 37 watts. All pain stimuli were delivered by Matlab which interfaced with the laser via a program built in Labview (Medical Physics department, Salford Royal NHS Foundation Trust). Participants submitted their intensity ratings of the pain using a keypad.

**Table 1:** Fluence. Energy density values expressed as fluence for each pain level.

| Energy density in J/cm<sup>2</sup> |  |  |  |
| --- | --- | --- | --- |
|  | Low | Medium | High |
| Mean | 0.2 | 0.32 | 0.47 |
| Minimum | 0.14 | 0.18 | 0.13 |
| Maximum | 0.31 | 0.46 | 0.75 |

### 2.4. Procedure

Participants underwent two blocks: the laser and the electric block. These two blocks differed only in the instrument used in stimulus administration, the location of the stimulation, and the fact participants wore safety goggles during the laser block of the experiment. Blocks were counterbalanced across participants. Stimulus timing was kept consistent between the two blocks. In the laser block, fibre laser stimuli were delivered to the dorsum of the participant’s right forearm. A 3cm by 4cm grid was drawn on the arm before beginning the study, and the laser stimulus was aimed at a new box in every trial, to reduce sensitisation or skin damage from the laser heat. In the electric block, electric stimuli were delivered to the middle phalanx of the right index finger.

Before applying the EEG cap, participants underwent a psychophysics procedure to determine their subjective response to increasing stimulus intensities, separately for the laser and electric stimulation. The order was counterbalanced between participants. The first stimulus was very low, generally an intensity which would be below the threshold for pain in most people, and the intensity of the stimuli increased in a ramping procedure. We used a 0-10 VAS^1^ for the pain response, where a level 3 was when the stimuli became painful, and level 7 was at the point where the participant did not wish to experience a higher level of stimulation in the experimental session. Consequently, we attained a ‘low’ pain level 3, a ‘medium’ pain level 5, and a ‘high’ pain level 7. We repeated this procedure three times to determine the average intensities corresponding to these intensity ratings. We then ran a procedure where participants identified the intensity (low, medium or high) of 18 pulses to ensure that they were experiencing the pain stimuli as they had rated in the psychophysics procedure. If participants could not identify the intensity of at least 75% of the pulses, the ramping procedure was repeated until they performed above 75%.

Trials were pseudorandomised. On each experimental trial, participants viewed a fixation cross for 500-750ms, and then a veridical probability cue which signalled the outcome likelihood and magnitude for that trial for 500ms. The cue, presented for 500ms, either signalled a 75% likelihood of high pain and a 25% likelihood of medium pain, or a 75% likelihood of low pain and a 25% likelihood of medium pain. This was followed by presentation of a blank screen for 1500ms, and then the stimulus was delivered, with the blank screen continuing for 1000ms. A screen was then presented which prompted participants to numerically rate their subjective pain intensity on a visual analogue scale (VAS) using a keypad. Inter-stimulus interval was maintained at a 10 second minimum. See figure 1 for a timeline for each trial. Over the experimental session, the 75% likelihood of high pain cue was followed by 90 high pain intensity stimuli and 30 medium pain intensity stimuli. The 75% likelihood of low pain cue was followed by 90 low pain intensity stimuli and 30 medium pain intensity stimuli. This meant on the relatively rare trials in which a medium pain intensity stimulus was administered, participants were expecting a high probability of either high or low pain, depending on the cue, which was crucial to our research question. At the end of every 7 minute block, participants rated the unpleasantness of the high, medium and low pain, on a 10-point VAS, following previous research (Pascalis, Chiaradia, & Carotenuto, 2002). In a subset of trials we presented a cue signalling 100% likelihood of receiving a medium pain, but these trials were not included in the analysis. We did not analyse these trials as they were relevant to a different analysis. Participants had a 2 minute break every 7 minute block.

**Figure 1:**
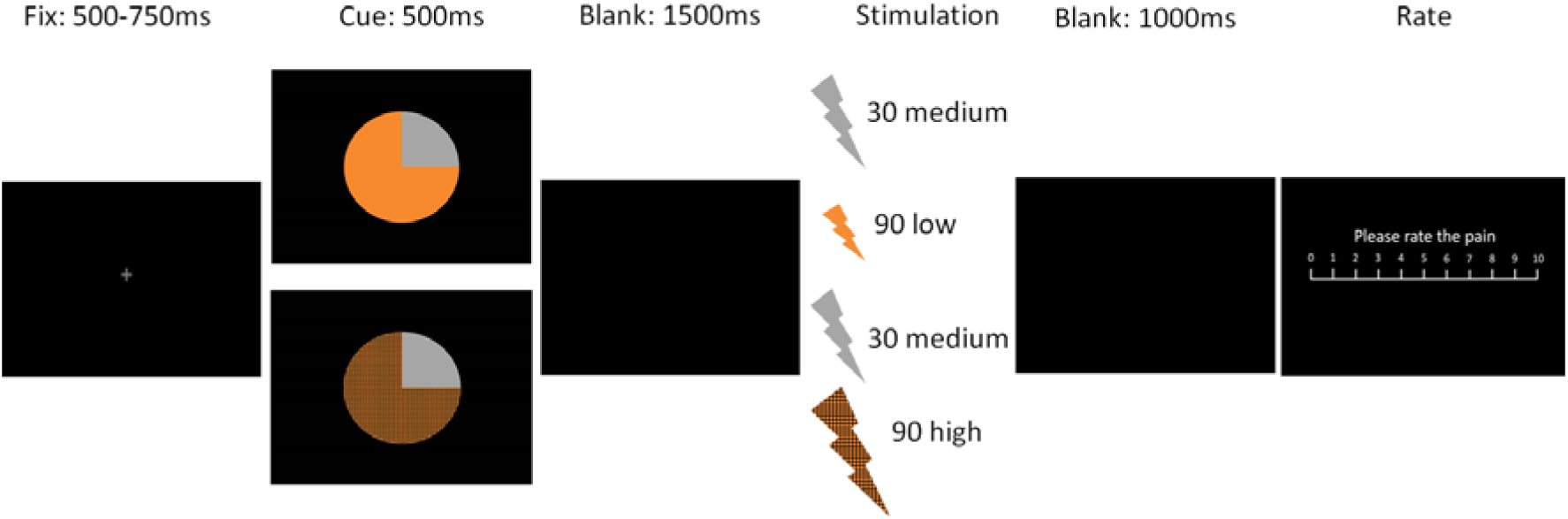
Trial timeline. Participants viewed a fixation cross for 500-750ms, followed by a veridical probability cue signalling the probability of receiving low (light orange) or medium (grey) pain in the low pain expectation trials, and viewed the same cue signalling the probability of receiving high (dark orange) or medium pain (grey) in the high pain expectation trials, for 500ms. After presentation of a blank screen for 1500ms, the painful stimulus was delivered, followed by presentation of a blank screen for 1000ms. Finally, participants were prompted to rate the VAS of the painful stimulus using a keypad.

### 2.5. EEG recording

Continuous EEG recording was acquired at a sampling rate of 1000 Hertz (Hz) using a 64 electrode Active-Two amplifier system (Biosemi, Amsterdam, Netherlands) with Biosemi acquisition software (BioSemi, Netherlands). An active and passive electrode replaces the ground electrode to create a feedback loop that drives the average potential of the subject (the common mode voltage) as close as possible to the analogue-to-digital converter reference voltage in the analogue-to-digital box. Impedances were kept at 20 KΩ or less. The experiment was conducted in a quiet room.

### 2.6. Behavioural data analysis

We firstly aimed to test for any psychophysical differences between the laser and electric pain stimulation. We conducted a t-test on the number of 5% stimulation intensity increases required to reach a VAS level seven pain rating per participant between laser and electric pain in the psychophysics procedure. A significant difference would indicate a difference in the increase of perceived pain per 5% increase in stimulation intensity between the two conditions. We also used Pearson’s correlation to test for correlations between average VAS rating and stimulation intensity across participants, where a significant correlation would indicate a consistent relationship between a 5% increase in intensity and a corresponding increase in VAS rating.

Next we conducted two 2 (stimulat or: laser/electric) × 2 (pain intensity: high/low) within-subjects ANOVA, one for the pain intensity ratings and one for the pain unpleasantness ratings. Additionally, we conducted a 2 (stimulator: laser/electric) × 2 (pain expectation: high/low) within-subjects ANOVA for the pain intensity ratings of medium pain trials. Interactions were followed up with post-hoc Bonferroni-corrected t-tests. Finally, we conducted a linear mixed model in each laser and electric pain condition separately, with trial as a predictor variable, to test for any effect of time on VAS rating. Participant was treated as a random effect.

### 2.7. EEG data analyses

Four participants were removed from the analysis because there was not a clear N2-P2 laser component, defined as a negative trough followed by a positive peak within 150-1000ms post-stimulus. One participant withdrew from the study, and one participant was detected as an outlier using Tukey’s method of outlier detection, which holds the advantage that it does not depend upon a mean or standard deviation, and so is resistant t o extreme values in the data (Tukey, 1977). This participant was removed from the data. The EEG signal for the remaining 14 participants was preprocessed using SPM12 (Ashburner et al., 2013; Litvak et al., 2011). Separate pre-processing pipelines were carried out for the pre-stimulus SPN and post-stimulus P2. Extracted data were analysed using SPSS.

#### SPN

The signal was referenced to the mean of all scalp electrodes, downsampled to 200 Hz and filtered with a lowpass Butterworth filter (30 Hz). Data were not highpass filtered as this could remove low frequency slow-wave anticipatory potentials. Epochs were extracted 200ms before the presentation of the probability cue to 1000ms after delivery of the pain stimulus. An absence of a highpass filtering step could introduce noise into the data and lead to an unnecessarily high artefact rejection rate. To avoid this, artefact rejection with a threshold of 60uV was applied to highpass (0.1Hz) filtered data to create a list of artefacts in the data. This artefact information was then applied to the actual data. In the laser pain conditions, the following trial numbers remained across participants: Cue low get low intensity pain (M=70.4, SD=13.5), cue low get medium intensity pain (M=24.9, SD=4), cue high get high intensity pain (M=66.9, SD=13.9), cue high get medium intensity pain (M=23, SD=4.2). In the electric pain conditions, the following trial numbers remained across participants: Cue low get low intensity pain (M=70, SD=11.8), cue low get medium intensity pain (M=23.4, SD=4.1), cue high get high intensity pain (M=66.5, SD=14.4), cue high get medium intensity pain (M=24, SD=3.3). Averaged across conditions, the difference in number of remaining trials between laser versus electric pain conditions was less than one trial. Accordingly, we do not anticipate any influence of trial number on results. Single-trial data were averaged separately for the eight conditions using the “robust averaging” method in SPM12b (Litvak et al., 2010). Based on previous research, data were extracted from the time-window 500ms to 0ms before pain delivery (Brown, Seymour, Boyle, et al., 2008a). We ran a 2 (stimulator: laser/electric) × 2 (pain expectation: high/low intensity) within-subjects on these data. We also used Pearson’s correlation to test for correlations between the average SPN and EEP amplitude, VAS and unpleasantness ratings across participants. Finally, we conducted a linear mixed model in each laser and electric pain condition separately, with trial as a predictor variable, to test for any effect of time on SPN amplitude. Participant was treated as a random effect.

#### P2

The re-referenced, downsampled signal was filtered with a Butterworth filter between 0.5 and 30 Hz. Epochs were extracted 700ms before the pain delivery to 1000ms after. Data underwent artefact rejection at a threshold of 60uV. In the laser pain conditions, the following trial numbers remained across participants: Cue low get low intensity pain (M=74.2, SD=15.8), cue low get medium intensity pain (M=24.4, SD=4.2), cue high get high intensity pain (M=66.1, SD=15), cue high get medium intensity pain (M=23.1, SD=5.2). In the electric pain conditions, the following trial numbers remained across participants: Cue low get low intensity pain (M=71.5, SD=12.3), cue low get medium intensity pain (M=23.6, SD=4.1), cue high get high intensity pain (M=68.3, SD=11.8), cue high get medium intensity pain (M=22.9, SD=5.4). Averaged across conditions, the difference in number of remaining trials between laser versus electric pain conditions was less than one trial. Accordingly, we do not anticipate any influence of trial number on results. Single-trial data were averaged separately for the eight conditions using the “robust averaging” method in SPM12b (Litvak et al., 2010), and filtered with a lowpass Butterworth filter (30Hz) to remove high frequency noise introduced by robust averaging, in order to use individual average data files to identify pain-evoked P2 latencies and topographies. The latency of 50% of the maximum amplitude of the grand average across all conditions and participants was identified (Luck, 2005). This was 267-450ms post-stimulus for the EEP, and 410-645ms for the LEP. Though the LEP latencies are late, similar latencies have been reported in previous studies in response to both CO_2_ (Brown, El-Deredy, & Jones, 2014) and thulium laser stimulation (Almarzouki, Brown, Brown, Leung, & Jones, 2017). Each participant’s maximum within this latency was identified, and ±40ms window was extracted around this, to include the peak of the maximum. See figure 2 for an illustration of the electrodes extracted in the analysis. The average amplitudes across this time window in each condition were analysed. We conducted two within-subjects ANOVAs to examine the extracted data, one with the factors 2 (stimulator: laser/electric) × 2 (pain intensity: high/low), and one with the factors 2 (stimulator: laser/electric) × 2 (pain expectation: high/low intensity). Finally, we used Pearson’s correlation to test for correlations between the average P2 and VAS rating across participants. Finally, we conducted a linear mixed model in each laser and electric pain condition separately, with trial as a predictor variable, to test for any effect of time on ERP amplitude. Participant was treated as a random effect.

**Figure 2:**
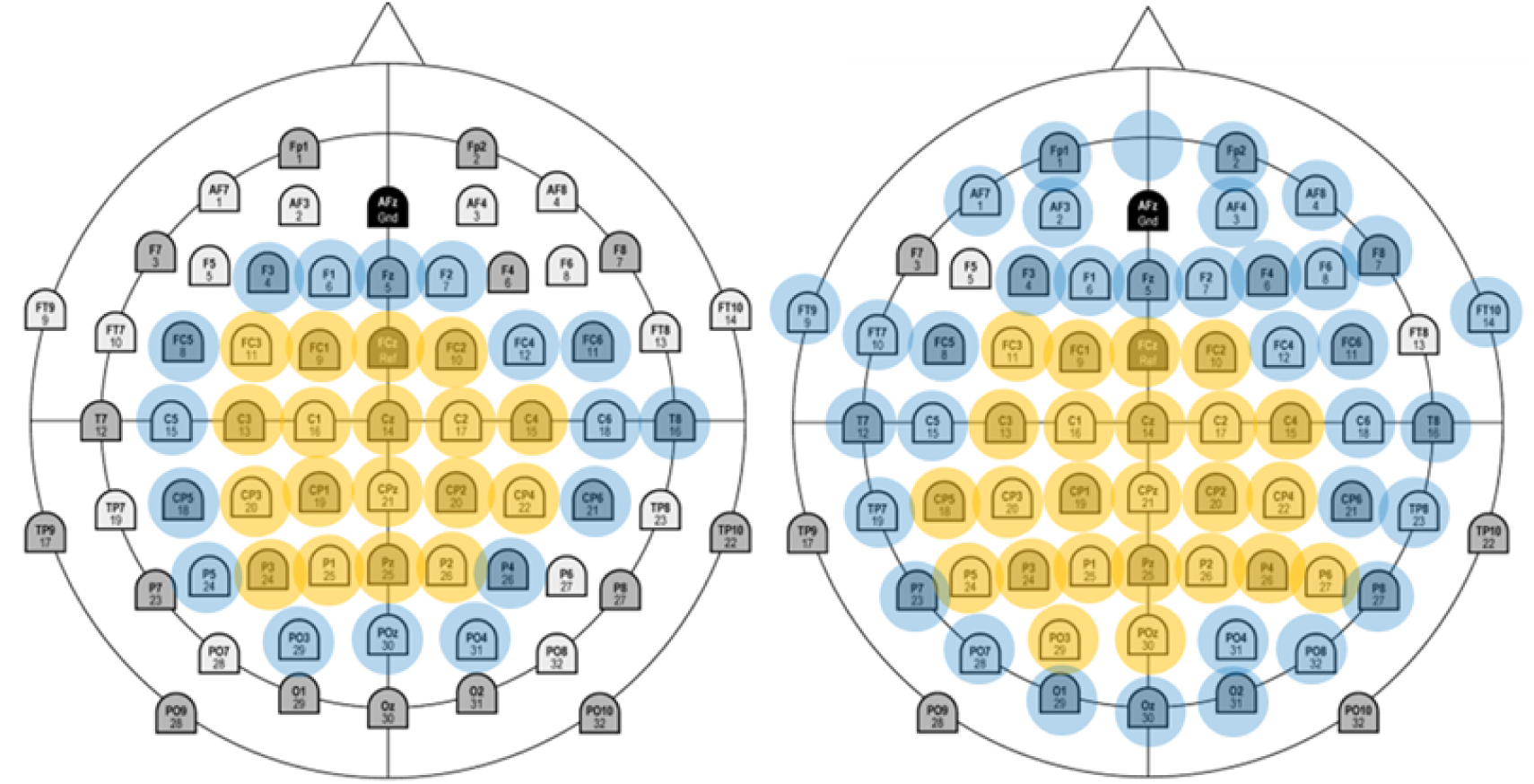
Electrodes. Electrodes expressing the P2 and selected for analysis for the electric (left) and laser condition. Electrodes extracted in the majority (at least 50%) of participants are highlighted yellow, and other electrodes extracted in blue. Across participants, the P2 peak was expressed at centroparietal electrodes, but more peripheral activation varied in topography between participants.

## 3. Results

### 3.1. Behavioural results

We first tested for any psychophysical differences between the laser and electric pain stimulation. In the psychophysics procedure, a paired samples t-test showed that across conditions, the number of 5% intensity increases from the lowest stimulation intensity to an intensity which elicited a level seven pain response was significantly higher in response to electric pain (M=8.33, SD=3.75) than to laser pain (M=5.4, SD=1.89) (*t*=-3.47, *p*=0.004, *d*=-1.085). The fact that a greater number of 5% intensity increases were required to elicit a level seven VAS rating to electric pain suggests that participants were more sensitive to increases in laser stimulation than electric stimulation. VAS rating significantly correlated with stimulator intensity in response to both laser pain (*r*=.82, *p*<0.001, *n*=87) and electric pain, though here the correlation coefficient was smaller (*r*=.49, *p*<0.001, *n*=124). The larger correlation coefficient in the laser pain condition further suggests that VAS ratings more closely reflected the 5% intensity increases in the laser pain condition compared to the electric pain condition.

We assessed the effects of pain intensity and stimulator on intensity rating. A 2 (stimulator: laser/electric) × 2 (pain intensity: high/low) within-subjects ANOVA revealed a main effect of stimulator (laser>electric) (f(1, 13)=6.813, 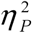 =.344) and a main effect of intensity (high>low pain) (f(1,13)=248.75, *p*<0.001, 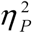 =.95), but no interaction (f(1,13)=.306, *p*=.59, 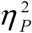 =.023), suggesting VAS score was overall higher in response to laser pain, but there was no difference between laser and electric pain in terms of how pain intensities were differentiated (see figure 4).

We next assessed the effects of pain cue and stimulator on intensity rating of the medium intensity pain trials. A 2 (stimulator: laser/electric) × 2 (pain expectation: high/low) within-subjects ANOVA revealed a main effect of stimulator (laser>electric) (f(1,13)=11.786, p=0.004, 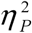 =.476) and a main effect of pain expectation (high>low pain cue) (f(1,13)=80.219, p<0.001, 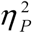 =.861), but no interaction (f(1,13)=78, p=.393, 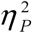 =.057), suggesting ratings were overall higher in response to laser pain, but that the high pain cue increased ratings equally compared to the low pain cue in both the laser and electric block (see figure 5).

We assessed the effects of pain intensity and stimulator on unpleasantness rating. A 2 (stimulator: laser/electric) × 2 (pain intensity: high/low) within-subjects ANOVA revealed a main effect of stimulator (laser>electric) (f(1,13)=25.268, *p*<0.001, 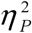 =.66) and a main effect of intensity (high>low pain) (f(1,13)=124.411, *p*<0.001, 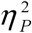 =.905), and an interaction (f(1,13)=13.686, *p*=0.003, 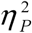 =.513). We executed post-hoc Bonferroni-corrected t-tests with a significance criterion of p<0.0125. They showed the effect of intensity to be significant in both the laser (*t*=12.933, *p*<0.001, *d*=3.74) and electric (*t*=7.429, *p*<0.001, *d*=1.934) condition, and the effect of instrument to be significant at high (*t*=5.668, *p*<0.001, *d*=1.33) but not low intensities at the significance threshold (*t*=2.821, *p*=0.014, *d*=0.83). In other words, high intensity laser and electric pain were rated as more unpleasant than low intensity laser and electric pain. High intensity laser pain was rated as more unpleasant than high intensity electric pain, but ratings for low intensity laser and electric pain did not differ.

Finally, we conducted a linear mixed model to test the effect of time on VAS rating within each condition, separately for laser and electric pain (table 2). In almost all conditions, there was a significant relationship between trial number and VAS rating. Beta values were positive across the laser pain conditions, indicating an increase in VAS ratings over time, and suggesting possible sensitisation. Beta values were negative in the electric pain conditions, indicating a decrease in VAS ratings over time, and suggesting possible habituation. In support of this, the average VAS rating across participants in significant trials increased from the first ten experimental trials to the final ten trials across laser conditions, and decreased from the first ten trials to the final ten trials across significant electric conditions (table 2).

**Table 2:** Effect of trial on VAS. Results of the linear mixed model, showing a positive relationship between laser VAS rating and trial, and a negative relationship between electric VAS rating and trial. Non-significant results are italicised.

| Stimulator type | Cued pain intensity | Delivered pain intensity | $R^2$ | $B$ | Standard error | $p$ | 95% Confidence interval | | Average (SD) VAS of first ten trials | Average (SD) VAS of final ten trials |
| --- | --- | --- | --- | --- | --- | --- | --- | --- | --- | --- |
| Laser | Low | Low | .05 | .004 | .004 | <.001 | .003 | .004 | 2.85 (1.07) | 3.59 (1.16) |
|  |  | Medium | .01 | .002 | .001 | .01 | .0004 | .003 | 4.46 (.81) | 4.72 (.71) |
|  | High | High | .01 | .001 | .0004 | <.001 | .001 | .002 | 6.56 (.91) | 6.79 (.87) |
|  |  | Medium | .02 | .002 | .001 | .003 | .001 | .004 | 5.82 (.91) | 6.19 (1) |
| Electric | Low | Low | .01 | -.01 | .003 | .01 | -.001 | -.0002 | 2.74 (.77) | 2.52 (.73) |
|  |  | Medium | .01 | -.001 | .01 | .05 | -.003 | .0002 | 3.94 (0.74) | 3.76 (1.04) |
|  | High | High | .04 | -.002 | .0003 | <.001 | -.003 | -.002 | 6.4 (.87) | 5.89 (1.49) |
|  |  | Medium | .04 | -.004 | .001 | <.001 | -.004 | -.002 | 5.33 (.93) | 4.83 (1.21) |

### Electrophysiological results

#### 3.2.1. Stimulus-preceding negativity

We subtracted the average response to the low pain expectation cue from the average response to the high pain expectation cue, and inspected the 500ms period preceding the pain stimulus in the average “difference” topography for each of the laser and electric stimulators. In the electric conditions we observed a negative difference across central-parietal electrodes which peaked at CP3 and CP5 (see figure 3), congruent with parietal anticipatory activity to electric pain shown in previous work (Berns et al., 2006; Hoflle et al., 2013; Lin et al., 2013). This was supported by a negative potential across conditions at these same electrodes. In the laser condition we observed a negative difference at C2 and Cz (see figure 3). We therefore extracted amplitudes from electrodes CP3 and CP5 for the electric condition, and C2 and Cz for the laser condition. A 2 (stimulator: laser/electric) × 2 (pain expectation: high/low) within-subjects ANOVA revealed a main effect of stimulator (electric > laser) (f(1,13)=16.79, p=0.001, 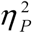 =.56), a main effect of cue (high > low) (f(1,13)=12.17, *p*=0.004, 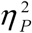 =.48), and an interaction (f(1,13)=5.92, *p*=0.03, 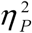 =.31) (see figure 3). Bonferroni-corrected paired samples t-tests with a significance criterion of *p*<0.025 showed the effect of cue to be significant in the laser (*t*=3.42, *p*=0.004, *d*=1.09) but not the electric pain condition (*t*=.81, *p*=0.43, *d*=0.22). These results suggest the amplitude of the electric SPN was overall more negative than the laser SPN, but the laser SPN differentiated pain intensity (cue low average amplitude was 4.41; cue high: 2.12), whilst although there was a numerical difference in the electric SPN, it was not significant (cue low average amplitude was −3.07; cue high: −3.4). SPN amplitude did not correlate with unpleasantness in either the laser (*p*=.84, *r*=-.041, *n*=28) or the electric condition (*p*=.61, *r*=-101, *n*=28), or with pain rating in either the laser (*p*=.26, *r*=-.152, *n*=56) or the electric condition (*p*=.08, *r*=-.24, *n*=56). The SPN did not correlate with ERP amplitude in either the laser (*p*=.33, *r*=-.133, *n*=56) or the electric condition (*p*=.89, *r*=.018, *n*=56).

**Figure 3:**
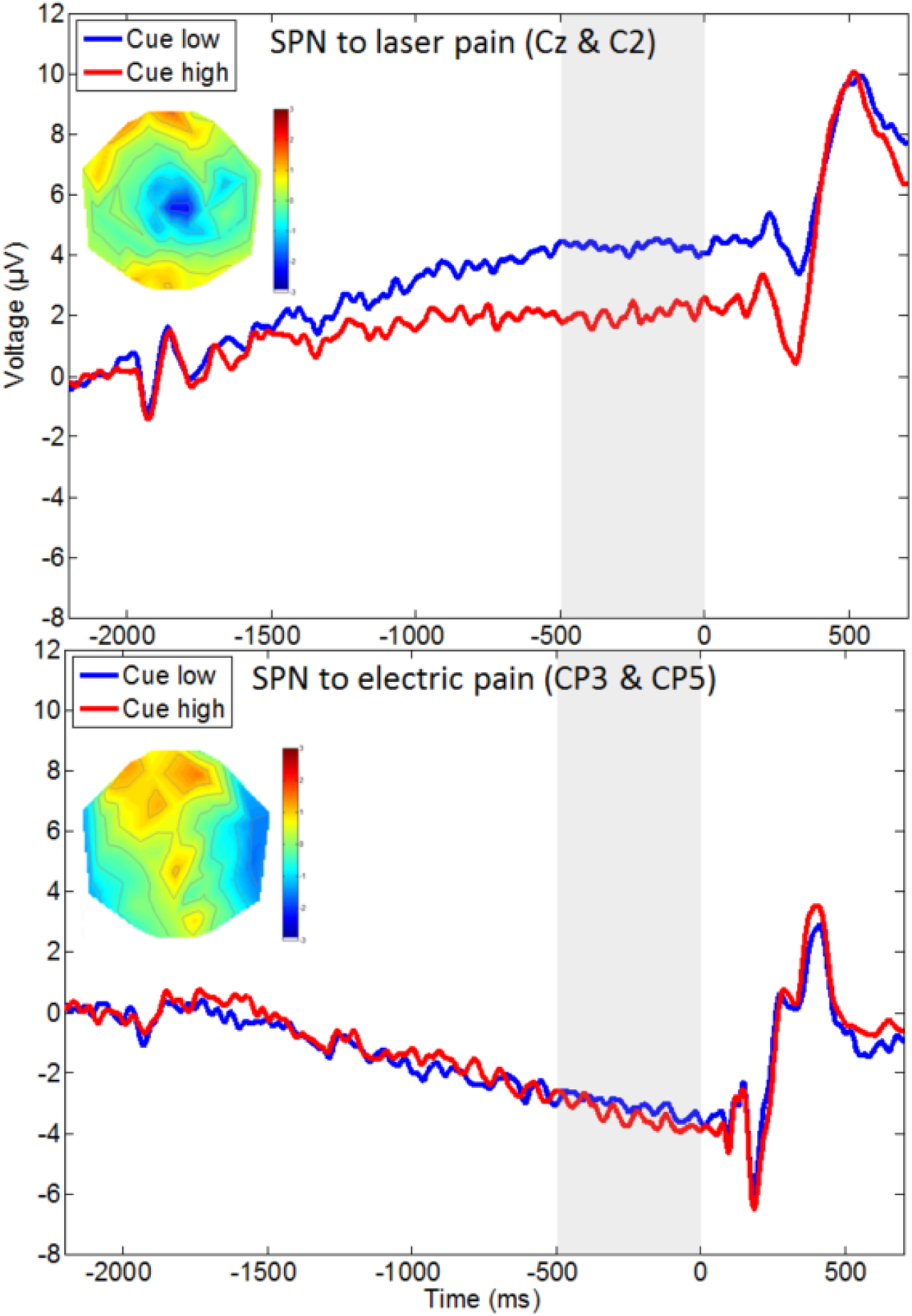
Anticipation. Stimulus-preceding negativity for high (red) and low (blue) pain cues for laser (upper plot) and electric (lower plot) pain. The probability cue was presented at −2000ms, and the pain stimulus delivered at 0ms. We analysed the signal from 500ms before pain stimulation (shaded grey). Topographic plots show topographies for the difference between high and low intensity pain expectation across the 500ms anticipatory period, scaled from −3 to 3 microvolts. Data were baseline-corrected to 200ms before presentation of the visual cue.

Finally, we conducted a linear mixed model to test the effect of trial on SPN amplitude within each condition, separately for laser and electric pain (table 3). Where results were significant in the laser pain condition, beta values were positive, suggesting SPN amplitude became less negative over time. In support of this, the average SPN amplitude over participants in the significant conditions decreased from the first ten experimental trials to the final ten trials (table 3).

**Table 3:** Effect of trial on SPN. Results of the linear mixed model, showing a positive relationship between SPN amplitude and trial. Non-significant results are italicised.

| Stimulator type | Cued pain intensity | Delivered pain intensity | $R^2$ | $B$ | SE | $p$ | 95% Confidence interval | | Average (SD) SPN amplitude of first ten trials | Average (SD) SPN amplitude of final ten trials |
| --- | --- | --- | --- | --- | --- | --- | --- | --- | --- | --- |
| Laser | Low | Low | .001 | .04 | .02 | .01 | .01 | .08 | -5.54 (5.47) | 3.03 (6.38) |
|  |  | Medium | .14 | .16 | .05 | <.001 | .07 | .25 | -17.12 (11.53) | -8.07 (5.88) |
|  | High | High | .001 | .03 | .02 | .09 | -.01 | .07 | 6.99 (7.83) | 23.66 (18.91) |
|  |  | Medium | .01 | .13 | .06 | .04 | .003 | .25 | 4.14 (6.88) | 5.66 (7.44) |
| Electric | Low | Low | -.0002 | .01 | .12 | .48 | -.02 | .03 | -3.88 (2.3) | -1.9 (4.91) |
|  |  | Medium | .0001 | .01 | .02 | .69 | -.03 | .04 | -3.74 (2.64) | -2.84 (2.48) |
|  | High | High | .002 | -.001 | .01 | .91 | -.02 | .02 | -4.15 (3.68) | -2.37 (4.04) |
|  |  | Medium | -.01 | .03 | .046 | .48 | -.06 | .12 | -4.07 (2.23) | -3.06 (3.73) |

### Effects of delivered pain intensity

Here we report ERP results from the veridically cued pain conditions, where a high pain intensity cue was followed by a high pain stimulus, and a low pain intensity cue was followed by a low pain stimulus. Peak latency of the EEP was shorter than LEP. The average peak latency across participants was 303ms (min=235ms, max=355ms, SD=43.3) to 383ms (min=325ms, max=455ms, SD=43.3) for EEPs, compared to 464ms (min=395ms, max=580ms, SD=55.3) to 544ms (min=475ms, max=660ms, SD=55.3) for LEPs. A 2 (stimulator: laser/electric) × 2 (pain intensity: high/low) within-subjects ANOVA on the voltages revealed a main effect of stimulator (electric>laser) (f(1,13)=24.69, p<0.001, 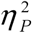 =.66) and a main effect of intensity (high>low pain) (f(1,13)=43.23, *p*<0.001, 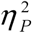 =.77), but no interaction, (f(1,13)=1.97, *p*=0.18, 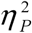 =.13) which indicates ERPs were higher in response to electric pain, but the effect of intensity was equal in response to both laser and electric pain (see figure 4). VAS ratings significantly correlated with ERP amplitude in the laser (*r*=.69, *p*<0.001, *n*=28) but not the electric condition (*r*=.31, *p*=.11, *n*=28).

**Figure 4:**
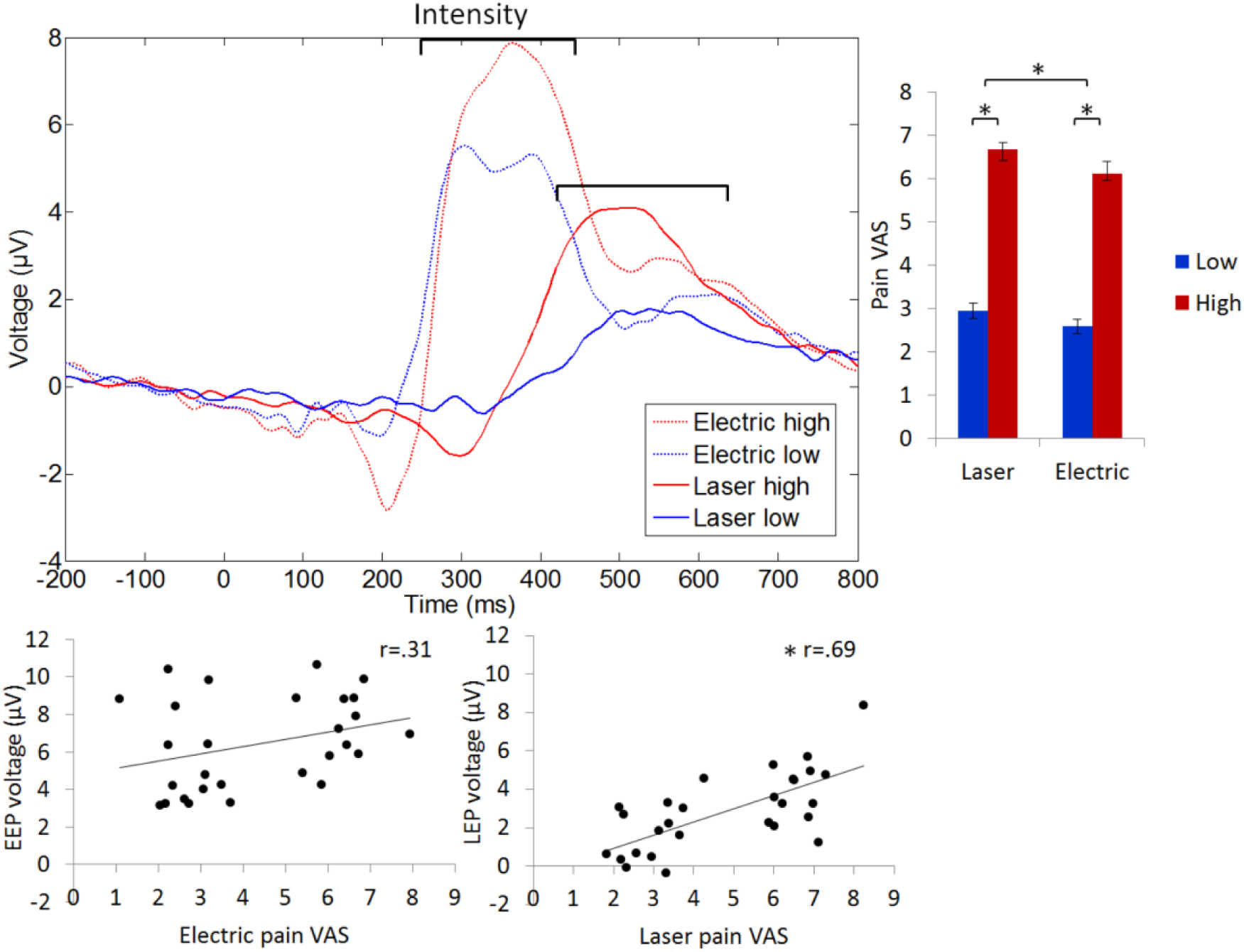
Intensity. Upper left panel: pain-evoked potentials for high (red) and low (blue) electric (dashed line) and laser (solid line) pain-evoked potentials at peak electrodes for each participant. The painful stimulus was delivered at 0ms. Bars show the window of analysis for each stimulator type. Upper right panel: average intensity VAS of the stimuli for the laser and electric pain. Red bars depict average VAS rating score for high intensity stimulation, and blue bars average VAS for low intensity stimulation. Error bars represent standard error of the mean. Lower panel: scatterplots showing correlations between EEP amplitude and VAS (left), and the LEP amplitude and VAS (right).

#### 3.2.2. Effects of expectations on medium pain stimuli

Here we report ERP results from the conditions where high and low pain intensity cues were followed by a medium intensity pain stimulus. A 2 (stimulator: laser/electric) × 2 (pain expectation: high/low) within-subjects ANOVA on the medium stimulus intensity pain-evoked potentials revealed a main effect of stimulator (electric>laser) (f(1,13)=15.6, *p*=0.002, 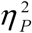 =.55) and a main effect of cue (high>low) (f(1,13)=5.71, *p*=0.033, 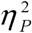 =.31), but no interaction (f(1,13)=084, *p*<0.78, 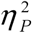 =.01), suggesting ERPs were higher in response to electric pain (as already noted in the section above), but the effect of cue was equal in response to both laser and electric pain (see figure 5). VAS rating did not correlate with ERP amplitude in either the laser (*p*=.26, *r*=.22, *n*=28) or the electric condition (*p*=.39, *r*=-.17, *n*=28).

**Figure 5:**
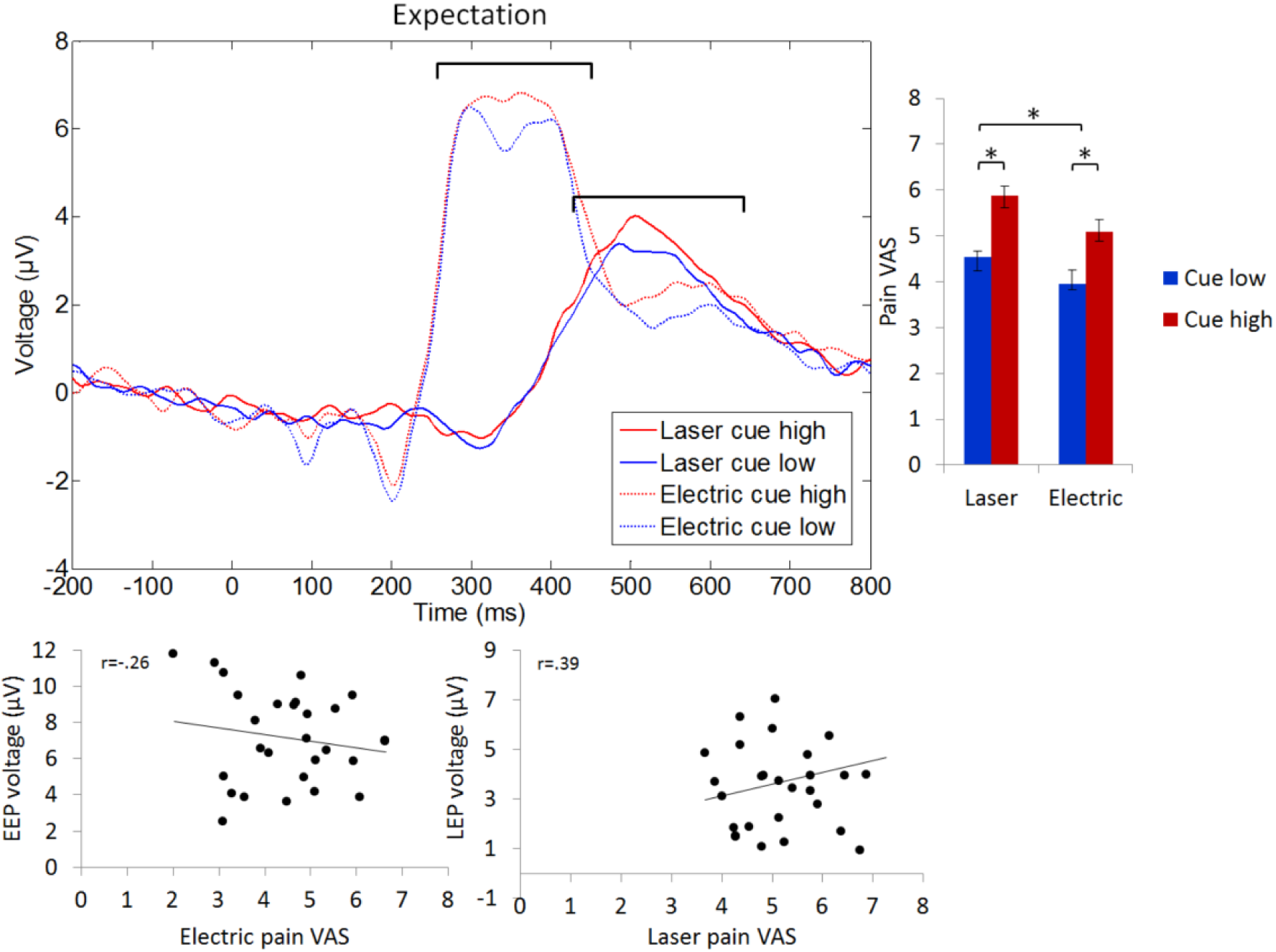
Pain expectation effects on LEP’s. Upper left panel: effect of high (red) and low (blue) pain cue on medium electric (dashed line) and laser (solid line) pain-evoked potentials at peak electrodes for each participant. The painful stimulus which in this case was always of medium intensity was delivered at 0ms with low and high intensity cues. Bars show the window of analysis for each stimulator type. Upper right panel: Average VAS intensity ratings of the stimuli for the laser and electric pain, high and low pain expectation. Error bars represent standard error of the mean. Lower panel: scatterplots showing the non-significant correlation between EEP amplitude and VAS (left), and LEP amplitude and VAS (right).

Finally, we conducted a linear mixed model to test the effect of trial on ERP amplitude within each condition, separately for laser and electric pain (table 4). In all conditions, there was a significant relationship between trial number and EEP amplitude. Betas were negative in almost all conditions, suggesting both LEP and EEP amplitude reduced over time. In support of this, the average ERP amplitude over participants decreased from the first ten experimental trials to the final ten trials (table 4).

**Table 4:** Effect of trial on ERP. Results of the linear mixed model, showing a negative relationship between LEP and EEP amplitude and trial.

| Stimulator type | Cued pain intensity | Delivered pain intensity | $R^2$ | $B$ | SE | $p$ | 95% Confidence interval | | Average (SD) ERP amplitude of first ten trials | Average (SD) ERP amplitude of final ten trials |
| --- | --- | --- | --- | --- | --- | --- | --- | --- | --- | --- |
| Laser | Low | Low | .01 | -.02 | .01 | <.001 | -.31 | -.01 | 2.57 (3.16) | .48 (2.04) |
|  |  | Medium | .06 | -.15 | .03 | <.001 | -.21 | -.09 | 3.97 (3.53) | 2.25 (1.7) |
|  | High | High | .01 | -.03 | .01 | <.001 | -.05 | -.02 | 3.93 (3.55) | 1.7 (3.25) |
|  |  | Medium | .04 | -.13 | .03 | <.001 | -.18 | -.06 | 4.48 (2.85) | 3.06 (2.69) |
| Electric | Low | Low | .01 | -.02 | .01 | <.001 | -.03 | -.01 | 6.71 (3.08) | 5.68 (2.5) |
|  |  | Medium | .03 | -.09 | .03 | <.001 | -.14 | -.04 | 7.61 (2.92) | 6.34 (2.94) |
|  | High | High | -.02 | -.02 | .01 | <.001 | -.03 | -.01 | 7.83 (2.49) | 6.91 (2.34) |
|  |  | Medium | .11 | -.13 | .02 | <.001 | -.17 | -.08 | 8.09 (2.95) | 6.53 (2.45) |

## 4. Discussion

Our results show that electric and laser stimulation elicits ERPs of different latencies and amplitudes, but these ERPs, and the corresponding subjective intensity ratings, were equally modulated by cue-evoked expectations. However, the intensity of the pain experience only correlated with the LEP and not the EEP. We also show morphological and topographical differences in the anticipatory SPN between the two stimulation types. Although the SPN is expressed in response to both stimulation types, it only differentiates intensity expectation significantly in response to the laser but not the electric pain stimulus.

### 4.2. Methodological discussion

The EEP, the marker for electric pain, differed in several ways to that for laser pain LEP. These differences can be explained by the fact that electric stimulation activates large myelinated somatosensory Aβ fibres as well as nociceptive Aδ fibres. Firstly, EEPs showed much shorter latencies than LEPs, consistent with previous studies comparing intracutaneous electrical stimulation with laser stimulation (Babiloni et al., 2007; Inui, Tran, Hoshiyama, & Kakigi, 2002). This suggests the method we used, transcutaneous electric stimulation, elicits potentials to a similar latency to more invasive intracutaneous stimulation. The reduced latency of the EEP can be related to the fact that Aβ fibres have a faster conduction velocity of 69 metres per second, compared to a conduction velocity of 10 metres per second in Aδ fibres (Tran, Lam, Hoshiyama, & Kakigi, 2001). Secondly, EEPs were of a greater amplitude than LEPs, in line with intracutaneous electric pain research, because Aβ fibre stimulation activates a larger number of thalamo-cortical units (Babiloni et al., 2007; Garcia-larrea et al., 2003; Gingold et al., 1991; Treede et al., 1999). This effect emerged despite the laser pain being rated as more intense and more unpleasant than EEPs. Related to this, when pain intensity was fully predicted by the cue, EEP amplitude did not correlate with intensity ratings, whereas LEP amplitude did. LEPs have been repeatedly shown to reflect intensity rating (Beydoun, Morrow, Shen, & Casey, 1993; Iannetti, Zambreanu, Cruccu, & Tracey, 2005; Ohara, Crone, Weiss, Treede, & Lenz, 2004), however, as EEPs reflect somatosensory Aβ fibre activity alongside Aδ fibre nociceptive activity, it is unsurprising that they did not correlate with perceived pain intensity alone, as the ‘noise’ of the unrelated somatosensory related activity would prevent identification of a relationship; this replicates previous EEP research (Rütgen et al., 2015). In summary, we show transcutaneous EEPs are earlier, higher in amplitude and do not correlate with pain intensity perception, in contrast to LEPs.

The laser pain stimulus was rated as more intense than the electric pain stimulus, despite participants undergoing a psychophysics procedure which was designed to calibrate stimulation to be equal between the two stimulator types. These differences in intensity rating are presumably due to changes between the psychophysics procedure and the main experiment, such as differing habituation rates and skin temperature changes associated with the two stimulators.

LEPs and EEPs were modulated equally by participants’ cued expectations. We propose that it is feasible to use either laser or electric pain to study cued expectation and pain processing. We observed an anticipatory SPN with differing topography and morphology between laser and electric pain, which differentiated pain intensity expectation during laser pain only. In this study, the SPN response to the intensity cue appears to be less sensitive prior to electrical stimulation than to laser stimulation. The differing topography, morphology and response to intensity of the SPN between conditions suggest they may originate from different neural generators.

We tested for effects of time on all measures of the pain response. Though significant results were small (see tables 2, 3 and 4), we did observe some differences between the two pain stimulation conditions. Laser pain perception increased over time, suggesting sensitisation to the stimulation, whereas electric pain perception reduced over time, suggesting habituation. ERPs in both conditions reduced over time, suggesting ERP habituation to the pain stimulus. SPM amplitude reduced over time in the laser but not the electric condition.

### 4.3. Theoretical discussion

Comparing evoked potentials and behavioural responses from the two stimulators alongside one another allows identification of any differences between the two stimulation types. LEPs and EEPs were modulated equally by participants’ cued expectation of the imminent intensity of pain, despite the higher subjective unpleasantness of the laser. Why did the higher affective impact of the laser pain not interact with the expectation cue? Emotion plays a significant role in pain; negative emotion and catastrophising increase perception of pain unpleasantness and intensity (Lin et al., 2013; Rainville, Bao, & Chrétien, 2005; Schupp, Berbaum, Berbaum, & Lang, 2005; Sullivan, Rouse, Bishop, & Johnston, 1997). The role of emotion in pain expectation is less clear, although studies suggest emotion plays a relevant role in expectation. For example, belief about the emotional impact of pain and confidence in the cue predicts the effect of expectation on pain (Brown, Seymour, El-Deredy, et al., 2008). Increased catastrophising also leads to higher engagement even with inaccurate cues, possibly due to the increased threat status of the cue in high catastrophisers (Van Damme, Crombez, & Eccleston, 2002). Placebo analgesia itself has been modelled as a reduction in pain-related anxiety (Flaten, Aslaksen, Lyby, & Bjørkedal, 2011; Morton et al., 2009). In our study expectations of pain intensity may have modulated the sensory-discriminative rather than the affective-motivational dimension of pain. Future work could manipulate the affective dimension of pain whilst maintaining a consistent intensity, to further disentangle these two closely related qualities. Analysis of the EEG sources, as in previous work, is likely to further clarify these issues (Brown, Seymour, Boyle, et al., 2008b).

The SPN expressed different topography and morphology between laser and electric pain, and differentiated between cue-evoked expectations of high and low laser but not electric pain. This finding adds to a somewhat inconsistent literature, where some studies show an SPN in response to laser but not electric pain, and others showing an SPN for electric pain (Babiloni et al., 2003, 2007; Seidel et al., 2015). The SPN for laser pain here differentiated between anticipation of high and low pain at central electrodes, as in previous studies showing sources in the anterior insula and cingulate in the laser pain SPN (Brown et al., 2008). However, the latter observations were only made for certain expectations and not for uncertain expectations. We observed an SPN to electric pain at centroparietal electrodes contralateral to the site of stimulation, in line with previous studies showing activity in the posterior insula and posterior cingulate during anticipation of electric pain (Berns et al., 2006; Hoflle et al., 2013; Lin et al., 2013). The contralateral topography of the electric SPN suggests activity of the lateral sensory-discriminatory somatosensory cortices, rather than a medial affective-motivational response.

The differing topographies of the SPN across stimulators were accompanied by a difference in morphology, with a greater negative amplitude for electric rather than laser pain overall. This is surprising, as the anticipatory SPN should increase for events with greater impact, here the subjectively higher intensity and more unpleasant laser pain. Interestingly, the SPN amplitude did not correlate with unpleasantness ratings, in either condition. Our results suggest the SPN may, under certain conditions, reflect processes related to somatosensory components of pain processing. However, as we did not manipulate this, we cannot draw firm conclusions about this proposed role of the SPN, but it is a key direction for future work.

There are some limitations to the comparison between laser and electric pain, because the two instruments require slightly different mode of stimulation, which could influence the results. First, the location of the laser stimulation was changed systematically between trials, but the location of the electric stimulation was kept constant. This is a variable which is inherent to laser pain studies (e.g. Lorenz et al., 2005; Morton et al., 2009; Watson et al., 2007). The fact that a new area of skin was stimulated on each laser trial could influence participant’s ability to discriminate the pain, although trial randomisation avoids any systematic bias associated with stimulation site and is therefore unlikely to affect the main results that we reported. The changes in the location of the laser stimulation also implied that it was less predictable than the electric stimulation. We aimed to minimise any unpredictability by moving the laser in a systematic and predictable pattern across the skin. Further, the laser was visible to the participant so they were able to predict the position of the laser. It is also worth noting that the two instruments stimulated different locations in the body which could introduce differences between the two conditions. However, in terms of cortical topographic representation, the change in location is relatively small (Bingel et al., 2004; Saladin, 2012; Stippich et al., 1999), so is unlikely to influence the pain-evoked potential. It is also of note that participants were not instructed to attend specifically to either the location or the unpleasantness of the pain, and so this may have been a source of variability between participants. Attention to either of these aspects of pain can influence the neural networks responding to pain and this may be why we did not see any correlation between SPN amplitude and unpleasantness or intensity ratings (Kulkarni et al., 2005). Further, while the use of 75/25% probability cues was justified in this study because the key research question was about pain modulation by positive and negative expectation, the relationship between SPN and ERP amplitude has been observed only in response to fully predictable pain intensity cues (Brown, Seymour, Boyle, et al., 2008b). Our results support these findings in that all cues contained an element of unpredictability and we did not observe a relationship between SPN and ERP amplitude. This indicates that future studies should be designed with a fully predictable cue in order to examine the relationship between anticipatory SPN and pain-evoked potential amplitude.

Responses to laser and electric pain were remarkably similar to one another, particularly in their modulation by cued expectation cues. When there is a reason to use many trials, our results suggest researchers should not hesitate to employ electric stimulation if expectation is the effect of interest. See table 2 for advantages of laser and electric stimulation. For future studies we recommend that the use of a 75/25 reinforcement schedule and a minimum of 30 trials recorded per condition are adequate to capture modulation of pain-evoked potentials by expectation in either laser or electric pain. Further research is required to test whether this effect can be captured across laser and electric pain under different pain expectation manipulations, for example in a placebo analgesia manipulation. Larger anticipatory effect sizes may be obtained when employing a ‘countdown’ anticipation period (e.g. Brown, Seymour, Boyle, et al., 2008b). We also recommend that, as here, future studies limit the maximum number of delivered fibre laser pain stimuli, to avoid any skin heating or sensitisation. Finally, generally in pain expectation studies, it would be prudent to maximise the predictability of the pain stimulation location as we have done here. This minimises any confounding effect of location-related surprise on intensity expectation effects.

We show that despite the absolute differences in intensity and unpleasantness ratings, and the latency and morphological differences of ERPs, intensity ratings and the marker for laser and electric pain (EEP and LEPs) are modulated equally by cue-evoked expectancies. Further studies are required to explore the potential of using the two techniques to access different aspects of the processing of pain anticipation. In view of the powerful effects of placebo and nocebo effects, which are substantially driven by negative and positive expectation (Amanzio et al., 2013; Atlas & Wager, 2012; Luana Colloca & Miller, 2011; Dodd, Dean, Vian, & Berk, 2017), both LEP and EEP methodologies have the potential to provide a physiological marker of individuals participating in clinical trials. EEPs provide a more practical method for larger-scale studies, and the results of this study provide motivation for exploring this further.

**Table 5:**
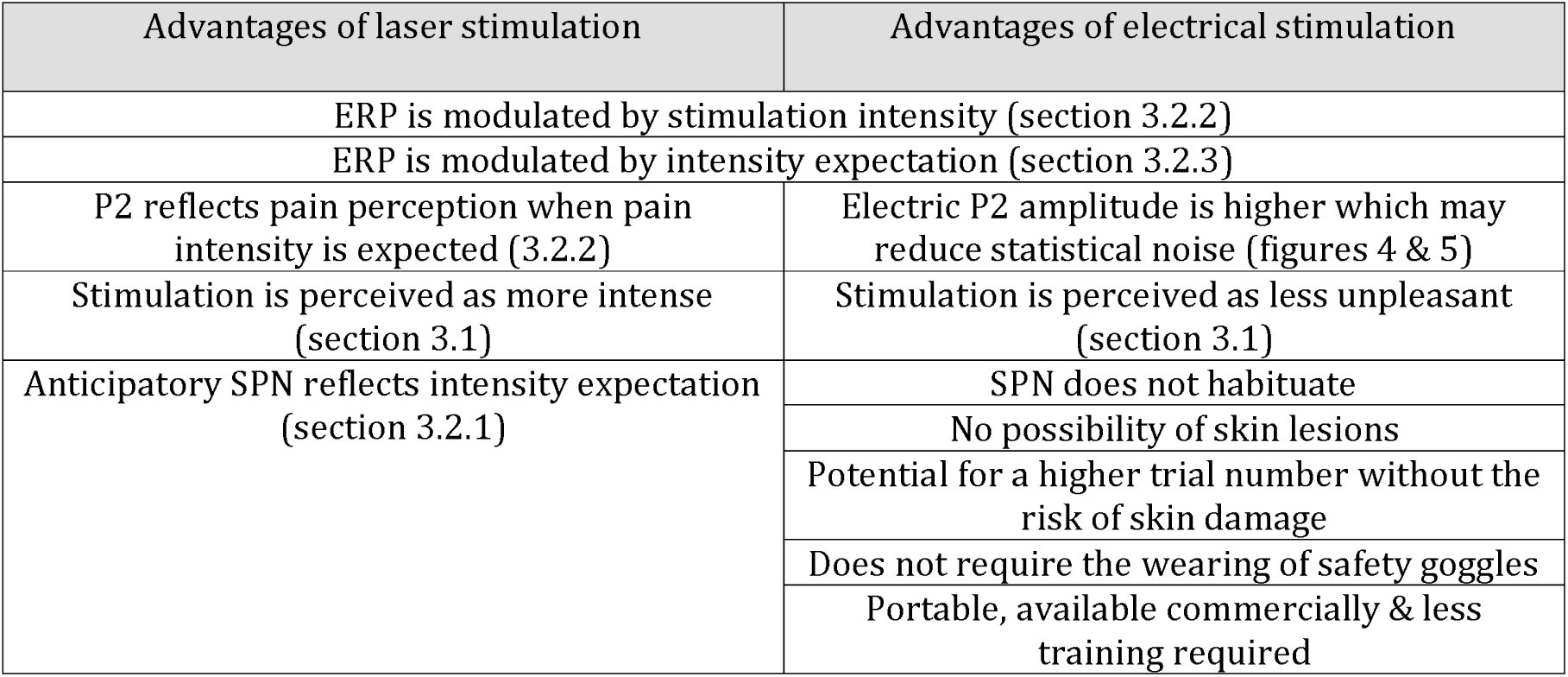
Laser versus electric stimulation. A comparison of advantages of laser and electric stimulation

## Acknowledgements

This work was supported by a studentship grant from the Medical Research Council, UK. We would like to acknowledge the Medical Physics team at Salford Royal Foundation Trust, especially P. Samraj and S. Watson, for their technical assistance in building the Labview program to operate the laser. We thank T. Rainey for his assistance with data collection and his technical expertise. WeD acknowledges the support of CONICYT, Chile, Basal project FB0008 and FONDECYT project 1161378. WeD is currently affiliated with the School of Biomedical Engineering, University of Valparaiso, Chile. The aforementioned sources of support did not have a role in study design, collection, analysis or interpretation of data, writing of the report, or the decision to submit the article for publication.

We declare no conflicts of interests.

## Footnotes

1 ERP: Event-Related Potential; LEP: Laser-Evoked Potential; EEP: Electric-Evoked Potential; fMRI: Functional Magnetic Resonance Imaging; rACC: Rostral Anterior Cingulate Cortex; SPN: Stimulus-Preceding Negativity; VAS: Visual Analogue Score

